# Network-Level Characterization of Spontaneous Calcium Activity in an In-Vitro Alzheimer’s Disease Model

**DOI:** 10.64898/2026.05.28.728474

**Authors:** Anna M. Emenheiser, Emma Gentry, Huijing Xue, Phillip Alvarez, Kate M. O’Neill, Kan Cao, Wolfgang Losert

## Abstract

The neurodegenerative disorder Alzheimer’s disease (AD) is widely known for biomarkers such as amyloid beta plaques and tauopathy, as well as functional differences in memory and cognitive ability. Despite this devastating functional impact, a large body of work only focuses on molecular biomarkers of AD. In this study, we investigate collective neural dynamics *in vitro* and assess how network-level properties differ between a well-established model of familial AD (FAD) and a newly developed *in vitro* accelerated model (acAD). The new model system reliably develops the key structural characteristics of AD in three weeks, but its calcium dynamics had not been characterized previously. Spontaneous network dynamics influences information processing as part of the internal network state. Here we measure this spontaneous activity of a network of hundreds of cells in each field of view. We find that the FAD model has a larger fraction of hyperactive cells, while the acAD model displays similar characteristics to healthy cells. Additionally, the FAD model has altered cooperation between cells, losing a proportion of highly correlated cellular activities, both for fast and slow coupling among cells. The acAD model is again consistent with healthy networks. Since the acAD model does not show the same spontaneous network dysfunction seen in FAD, it can enable measurements of changes in learning and memory associated with the plasticity, rather than the structure of the internal network state.

## Introduction

A recent report estimates that over 7.2 million adults in the United States had Alzheimer’s Disease (AD) as of 2025 (1). This is expected to grow to nearly 14 million by 2060 (1). The increasing prevalence of neurodegenerative diseases highlights the importance of understanding physiological changes that occur with aging. In order to better understand memory and cognition in healthy brains and the disease state, here we investigate collective neural dynamics *in vitro* to identify both established and previously unexplored network-level features in classical AD culture systems and a newly developed *in vitro* model of Alzheimer’s disease (AD). AD is pathologically characterized by amyloid beta (Aβ) plaques and tauopathy-related neurofibrillary tangles (1, 2). A large body of previous work has been dedicated to what is known as the amyloid hypothesis, focusing on the presence of Aβ plaques. However, recent studies are increasingly moving away from the amyloid hypothesis. There are instances where elderly individuals had significant Aβ plaques but were not diagnosed with AD (3, 4). Additionally, new research suggests that a lack of soluble Aβ_42_ may be more informative than the formation of plaques (5). Considering the discrepancies within the literature on Aβ, biomarkers unrelated to Aβ may be valuable to the field. While tau presents an appealing alternative, tauopathy can be found in other neurodegenerative diseases and does not recapitulate the Aβ plaques observed in AD (6).

Tauopathy and Aβ are not the only disruptions observed in AD. An alternative framework for AD explores characteristics of the disease through the lens of calcium dysregulation (7, 8). *In vitro* AD models have measured changes in calcium activity, specifically an increase in hyperactive neurons (9–11). Another study found a decrease in calcium signaling, though this was in the presence of T cells in the neural culture (12). In mouse models, it was found that neurons in close proximity to Aβ plaques were more likely to be hyperactive (13). Additionally, hyperactivity was shown in mice with mutations in presenilin 1, despite a lack of Aβ plaques (14). Patients with mild AD display neuronal hyperactivity, however, in late-stage AD, hypoactivity is observed due to neuronal damage and death (15). This was also demonstrated in human cell models, though further characterization is necessary due to a lack of a reliable AD cell model (15). We hypothesize that calcium dynamics across networks of *in vitro* neural networks will be affected by the progression of AD and may serve as a reliable biomarker. By measuring network dynamics, we avoid a hypothesis that is hyper-specific to one mechanism or pathway of AD.

When observing network dynamics *in vitro*, spontaneous activity, whether electrical or calcium spikes, is often considered “noise”. This is comparable to spikes recorded *in vivo* that are not the result of a target behavior or stimulus, or the variability between trials in neuronal responses (16). However, correlations between such signals can be considered for insights into the balance of excitatory and inhibitory neurons (17). New results also show that “noise correlations” provide an upper bound on the information representation of spatial information in the mouse brain (18). While noise correlations can be used to understand aspects of connectivity, they are not sufficient to understand the functional networks involved in information processing (19). The capability to process information and learn depends on the internal dynamic state of the network, which is made up of both the spontaneous activity and network plasticity (20). Thus studying cognitive disorders such as AD may benefit from separating these two contributors to the internal dynamic state of a neural network. Here we study spontaneous activity in multiple models of AD. While there are many existing *in vivo* mouse models of AD, potential treatments that have been taken to clinical trials have been largely unsuccessful (21–23). Additionally, previous *in vitro* models using human cell lines often require several months for the pathological hallmarks to present, increasing the difficulty of high-throughput testing for potential treatments (24).

The accelerated Alzheimer’s Disease (acAD) model utilizes three different familial AD (FAD) mutations: K670N/M671L (Swedish) and V717I (London) mutations (APPSL) and PSEN1 with the ΔE9 mutation (PSEN1ΔE9) (24). These FAD mutations are estimated to represent 15-25% of AD cases (25). APP and PSEN1 FAD mutations are known for complete penetrance, where all patients with either mutation will develop AD if they reach the typical life expectancy (25). The acAD model takes advantage of the rapid-aging disorder, Progeria (26). Progeria, or Hutchinson-Gilford progeria syndrome, is caused by a mutation to the lamin A gene, progerin (27). Progerin disrupts the scaffolding of the nuclear envelope (28). Although this mutation is not typically expressed in neural cells, by forcing the coexpression of Progeria and AD mutations, the neuronal network expresses AD pathological features more rapidly (within 3-4 weeks) (26).

Using human neural progenitor cells (hNPCs) as the basis for the model, the resulting self-assembled networks contain both neurons and astrocytes (26). Astrocytes directly participate in neural communication, through modulation of synapses and affecting information processing in neurons (29, 30). There is increasing evidence that astrocytes play an important role in learning and memory, specifically connected to behavioral differences (31–33). Astrocytes have also been implicated in AD directly, including observations of hyperactive calcium signaling (34–37). We believe the inclusion of astrocytes within *in vitro* models, such as those derived from hNPCs, is valuable for studying AD. This work aims to functionally characterize the spontaneous activity of this acAD model.

## Methods

### Lentivirus

Lentivirus was produced using HEK293T cells (ATCC) grown in DMEM + 10% FBS. Using Fugene 6 (Promega), the cells were co-transfected with psPAX2 (Addgene), pMD2.G (Addgene), and the lentiviral plasmid. At 48 hours and 72 hours, the media was collected and filtered using 0.45 µm filters and frozen at −80°C.

### Cell Culture

hNPCs from the ReNcell VM Human Neural Progenitor Cell Line (SCC008, EMD Millipore) were used. The undifferentiated hNPCs were cultured in proliferation media (ReN Maintenance Media with 20 ng/ml bFGF and 20 ng/ml EGF) and were maintained at 37 °C, with 5% CO_2_ and humidity control. Prior to seeding and differentiation, lentivirus carrying the mCherry or FAD plasmids was added for 48 hours in proliferation media with 8 µg/ml polybrene. The virus was removed and cells were further proliferated for 3 days. The hNPCs were plated on 35 mm glass-bottom dishes coated with Geltrex at a density of 3e5 cells per dish. The day after seeding, the media was changed to differentiation media (Brainphys with 2% 50X B-27 and 0.1% Heparin sodium), lacking bFGF and EGF growth factors to trigger differentiation.

Lentivirus carrying the GFP, GFP-Lamin A, or GFP-Progerin plasmids was added in differentiation media with 8 µg/ml polybrene for 48 hours, as represented in Fig 1A. Media was changed every 2-3 days to promote cell health until analysis at 14 or 21 days of differentiation. To control for the combination of the mutations for both diseases (FAD and Progerin) as well as the fluorescent tags included, ten different variations of genetic mutations, including one non-transduced control (CTRL), were tested S1. For brevity, only three cell lines are described in the results: CTRL, FAD, and acAD. The genetic constructs used are included in the supplemental materials S1.

**Fig. 1.**
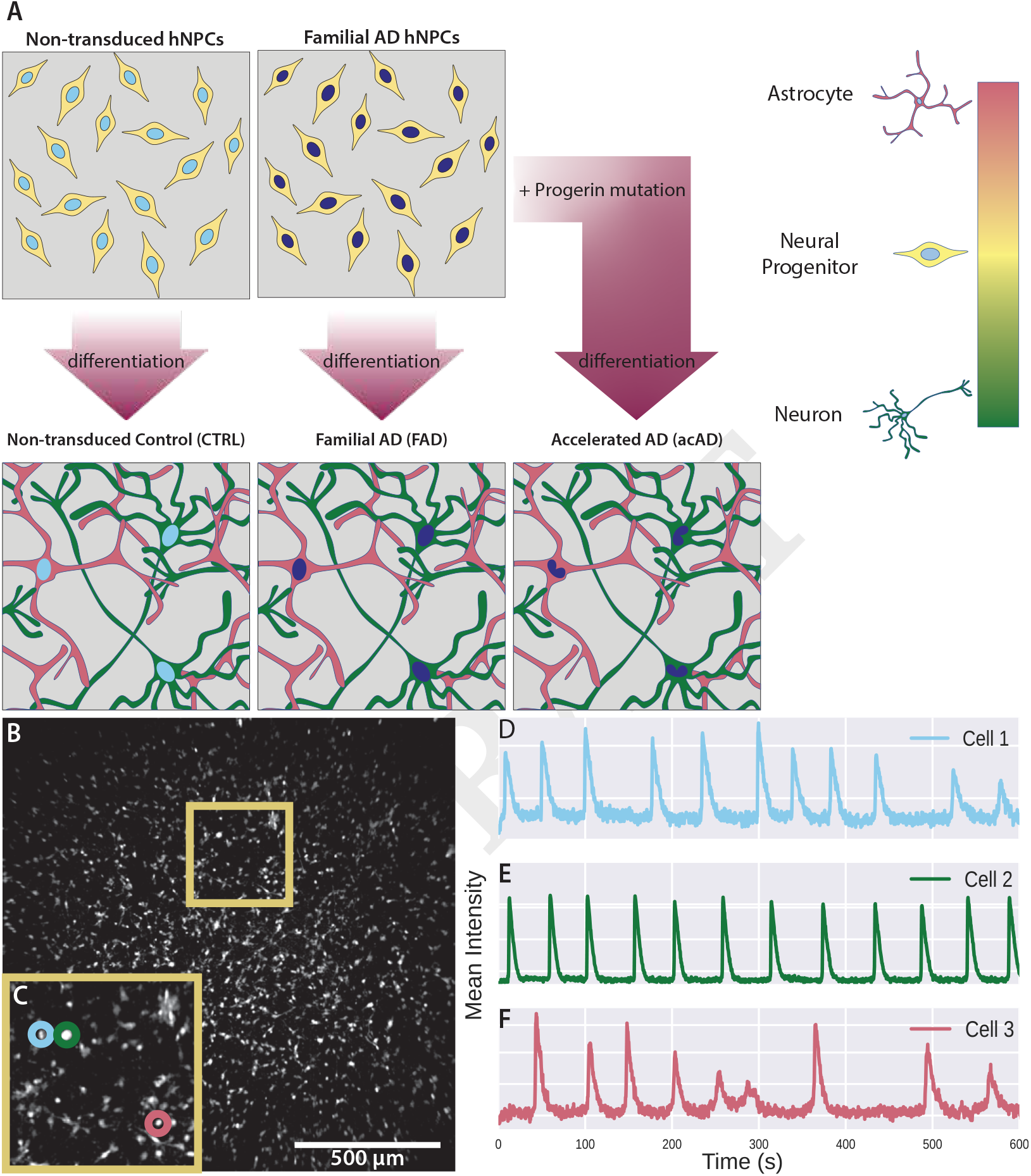
Overview of cellular models and typical imaging field of view. **A)** Schematic representation of the generation of the cellular models used. We begin with undifferentiated neural progenitor cells that self-assemble into networks of astrocytes and neurons as they differentiate. The acAD model is created using a combination of familial AD and Progerin mutations. Progerin is a mutation in the Lamin A gene, which causes a deformation of the nucleus as represented here by a different nuclear shape in the differentiated network. **B)** A typical field of view during widefield epifluorescence with background subtraction for visualization purposes only. Background subtraction was not performed in the image analysis workflow.**C)** A magnified inlay from the full field of view image, the original location is noted with a yellow box. Three cells of interest are highlighted. **D-F)** Raw fluorescent traces in time of the three cells of interest.

### Imaging

Imaging was performed at two and three weeks after differentiation, in order to compare a baseline (week two) to activity after we expect the acAD model to reach the AD-like state (week three). In order to be succinct, only week three is discussed in the results. Cells were incubated for one hour with a fluorescent calcium indicator, Calbryte™ 590 AM (AAT Bioquest), at a 1 µM concentration. After a full media change to remove excess dye, the cells were imaged using Micro-Manager software (38) on a custom multiscale microscope, using a 10X objective (Nikon, CFI Super Fluor, NA 0.5) together with a tube lens demagnified by 0.5X (final effective magnification of 5x) to capture a field of view up to 2.9 mm by 2.4 mm (Fig 1B). Timelapses were captured with the widefield epifluorescence modality, using a 561 nm laser (OBIS, Coherent), for 10 minutes at 3 Hz, with a sCMOS camera (pco.edge 5.5, Excelitas), while maintaining the cells at 37 °C, with 5% CO_2_ and humidity control (Ibidi).

### Image Analysis

After image capture, analysis was performed with a custom Python script adapted from previous work in our lab (39). Cell segmentation and fluorescent trace extraction was performed by CaImAn. The fluorescence trace was then normalized to the baseline fluorescence, ΔF/F, example traces shown in Figure 1D-F. Peaks in the normalized ΔF/F calcium traces were identified through an established peak finding method in Python. The total count of spikes normalized to the length of the recording determined the overall activity of a particular cell. The time-delay cross-correlation between pairs of cells was calculated based on these ΔF/F traces and was determined for all pairs of cells in an observed network. The zero-delay cross-correlation was used to understand instantaneous or fast communication within the network. The maximum cross-correlation was used to understand slower communication within the network. However, to avoid artificially inflating the cross-correlation values by selecting the absolute maximum, the lags were limited based on the distance between the cells, and the maximum was selected from that smaller range. Each of these three metrics (zero-lag cross-correlation, distance-limited cross-correlation, and activity) form distributions for a network, representing a variety of cell behaviors. For each of these three distributions, we measured the first four moments (mean, variance, skew, kurtosis) and the standard deviation.

### Statistics and Replicates

Figures 3-5 represent at least n=3 distinct biological replicates, the complete documentation of biological and technical replicates can be found in S1. For group comparisons across different disease mutations of the combined distributions, the Kolmogorov–Smirnov test with a Holm-Bonferroni correction was used due to it’s ability to detect differences in the shape of distributions beyond just different means. For comparisons of per-experimental metrics, the Kruskal-Wallis test was used to test differences in the mean of each metric. Final significance between conditions was determined using post hoc Games-Howell tests, which do not assume equal variances and are robust to unequal sample sizes. This approach was applied consistently across the three different measurements of activity, zero-delay cross-correlation and distance-limited cross-correlation. All statistical tests were performed in Python using standard scientific libraries.

## Results

Two approaches were taken in order to understand the behavior of each cell line across replicates. We begin with distributions of a given measurement for each technical replicate (Fig 2A-D). In the first representation, we considered each cell as an individual unit, and all cells identified across all experimental replicates of the same cell line were combined into one distribution of a measurement, such as spiking activity (Fig 2E). The combined histogram is weighted such that all biological replicates contribute equally, regardless of differences in the number of technical replicates and cells within technical replicates. Second, we evaluated the networks as the smallest unit, and measured the mathematical moments of the distribution from each recording before comparing these values across cell lines (Fig 2F). The violin plot is not weighted, but technical replicates belonging to the same biological replicate are indicated by the same color on the interior of the dot. These different representations, while not entirely independent, describe different aspects of the rich dynamics in this dataset. This framework will be used for all three measurements examined.

**Fig. 2.**
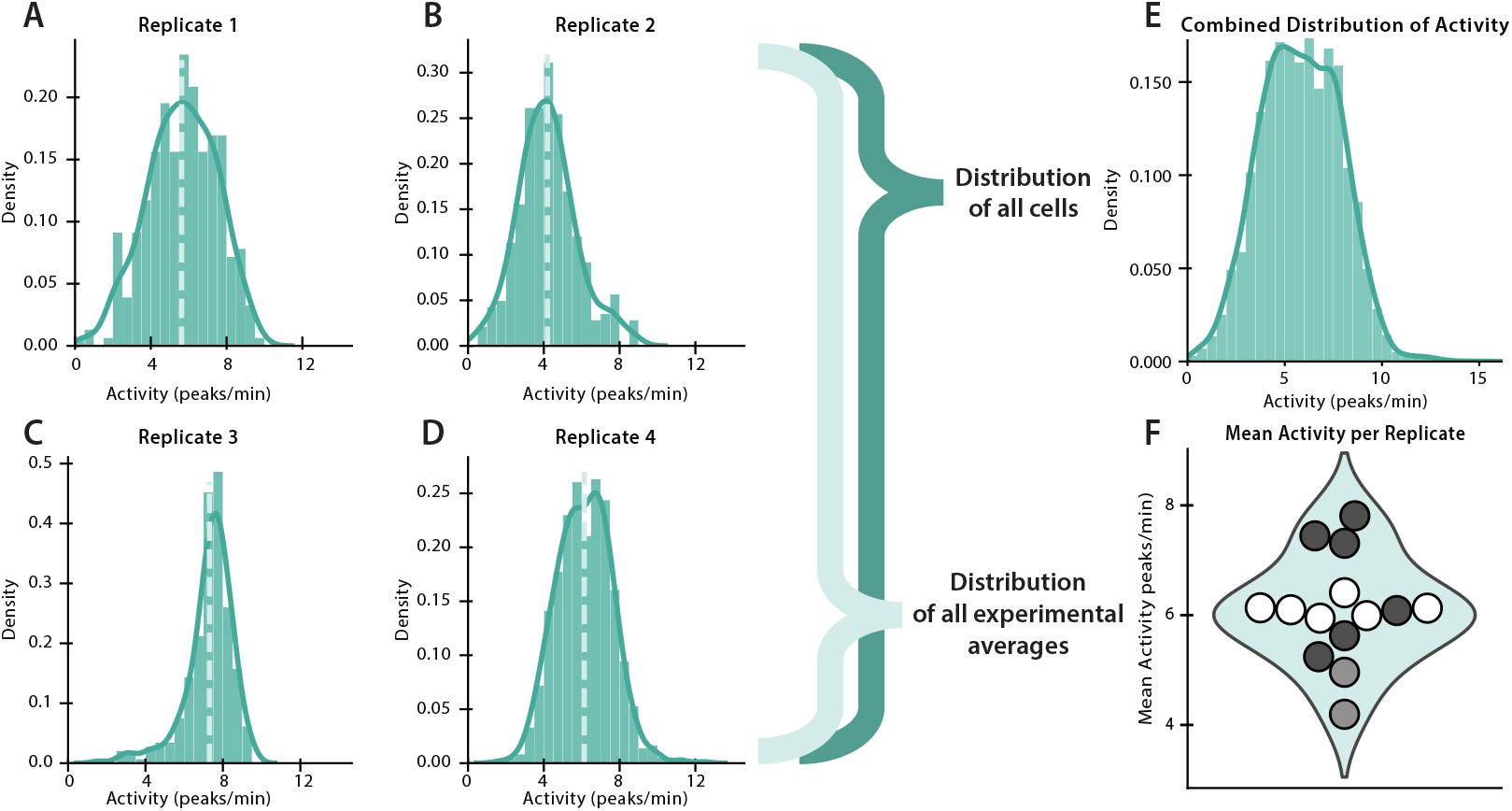
Schematic representation of distribution analysis workflow. **A-D)** Example density histograms of calcium activity (spikes/min) for all cells identified in each of four different experimental replicates. **E)** Combined density histogram of calcium activity for all cells identified across all experimental replicates. This is weighted such that each biological replicate contributes equally despite any differences in the number of technical replicates. **F)** Violin plot representing the means of each distribution of individual replicates. The color of each dot on the interior of the violin plot represents a different biological replicate. The overall plot is not weighted to account for this difference. Both representations of data are used for all three distributions measured: activity, zero-delay cross-correlation, and distance-limited cross-correlation.

### Ca^2+^ spike rate

To investigate if the disease mutations affected the spontaneous activity of the networks, we measured the Ca^2+^ spike rate of each cell. We found that despite the disease mutations in the FAD and acAD cell lines, none of the mutations resulted in a complete loss of activity (Fig 3 A-C). Kolmogorov–Smirnov (KS) tests of the combined distributions of individual cells (n = 21936 cells) indicated that all distributions are significantly different, although the effect size suggests only the FAD and acAD distributions deviate moderately (Table S2).

**Fig. 3.**
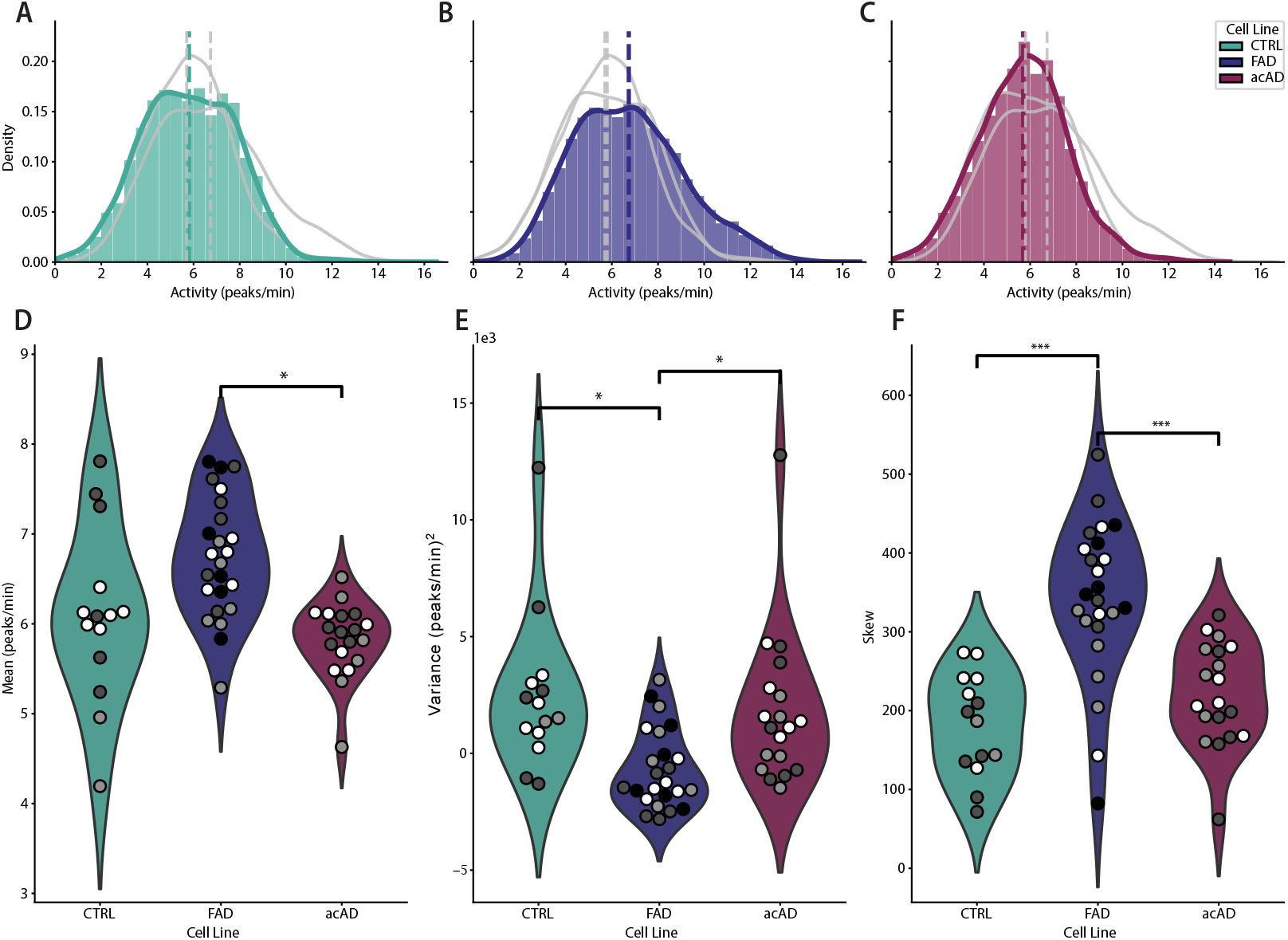
Characterization of calcium activity (spikes/min) of healthy and mutated cell lines. **A-C)** Density histograms of calcium activity (spikes/min) for all cells identified across all experimental replicates for CTRL, FAD, and acAD cell lines respectively. Dashed lines indicate the mean of each distribution. The main cell line is in color while the other two conditions are overlaid in grey. The histograms are weighted such that each biological replicate contributes equally despite any differences in the number of technical replicates.**D-F)** Violin plots of per experimental measurements of mean, variance, and skew (respectively) of the activity distribution. The color of each dot on the interior of the violin plot represents a different biological replicate. The overall plot is not weighted to account for this difference. Statistical significance is indicated by * (* p < 0.05, ** p < 0.01, *** p < 0.001).

Such differences persist in the network-level metrics, although to a different extent. A Kruskal-Wallis (KW) test of the per-experimental (n = 57 recordings) moments of the activity distribution revealed a significant effect of cell line on all three metrics (Table S5), suggesting substantial differences between networks. Post hoc Games-Howell tests confirmed a significant difference between the mean activity in FAD and acAD models (Fig 3D), all with large effect sizes. The variance and skew indicate further differences between the CTRL and FAD models (Fig 3E-F). Across all metrics, the FAD model is the most significantly different from CTRL and acAD networks. The differences in variance and skew describe a population of hyperactive cells with more calcium spikes over the period of observation. This can be seen in the combined distributions as well (Fig 3B). The acAD model on the other hand does not contain a population of hyperactive cells, maintaining a baseline level of activity similar to the CTRL.

### Time-delay Cross-Correlation

We explored not only single-cell behavior but also the functional connectivity between cells. Consistent with established methods for analyzing neural ensembles, we quantified correlation between all pairs of cells in the network. However, rather than using a Pearson correlation, we chose to use time-delay cross-correlation. Our networks consist of both neurons and astrocytes, so we used different time-delays in order to differentiate between fast and slow communications. Fast communication is captured by the zero-delay cross-correlation, which accounts for the near-instantaneous electrical signaling of neurons. Slower communication, such as astrocytic signaling or other non-electrical communication between neurons, is measured by the time-delayed cross-correlation. Rather than selecting the absolute maximum value of cross correlation and artificially inflating the distribution, we limited the time-delays that were allowed for each cell pair based on distance (further described in S5). We use the same framework as described above to interrogate the data. The zero-delay cross-correlation similarly displays significant differences between the combined distributions: KS tests of the cell-pair cross-correlation values (n = 5,545,197 cell pairs) revealed that all distributions are significantly different, although the effect sizes are only moderate (Table S3). Notably, there is a smaller second peak of high zero-delay cross-correlation coefficients in the healthy CTRL distribution (4A) that is not present in the other distributions (4B-C). This indicates a loss of cell-pairs with high instantaneous cross-correlation, potentially representing a loss of strong functional connections between neurons in the FAD and acAD models.

The network-level metrics indicate differences in the zero-delay cross-correlation but only in the variance and skew (KW (n = 57 recordings, Table S6). No pairwise differences in the mean reached statistical significance, but moderate to large effect sizes suggested consistent trends across groups that were consistent with those observed in the other metrics (Fig 4D). In contrast, the variance and skew of the zero-delay distributions are more significant (Fig 4E-F). Again, the FAD networks are the most different, confirming a disruption of functional connectivity at both the cell-pair and network levels. The smaller skews of FAD networks indicates a diminished tail in high cross-correlation pairs (Fig 4F). This is consistent with the trend visible in the combined distributions, and indicates that this loss of strong functional connections is true across all FAD networks. The CTRL and acAD networks have a wider spread in skew values, indicating higher biological variability in the functional connectivity and coordination.

**Fig. 4.**
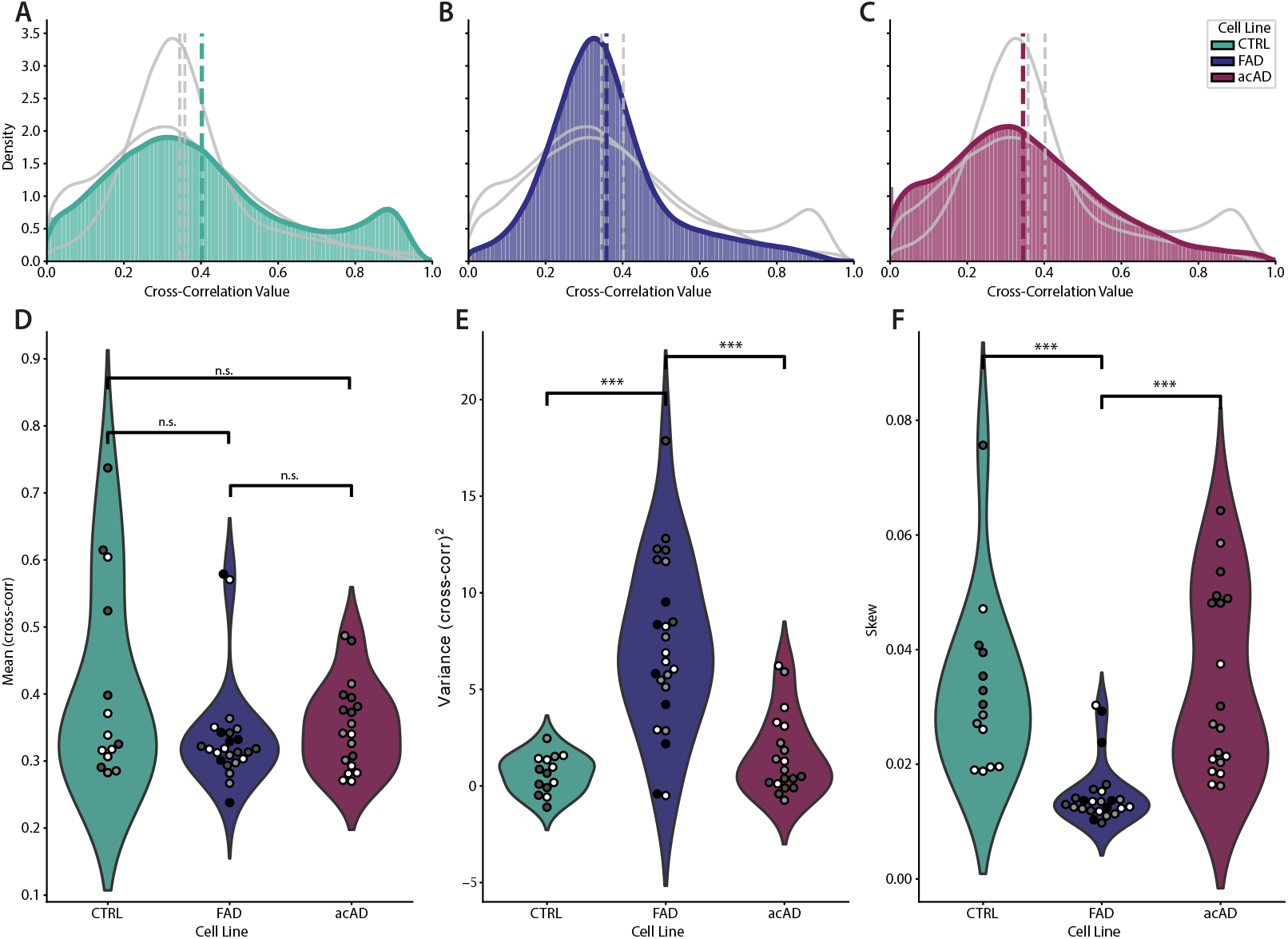
Characterization of zero-delay cross-correlation distributions of healthy and mutated cell lines. **A-C)** Density histograms of zero-delay cross-correlation coefficients for all cells identified across all experimental replicates for CTRL, FAD, and acAD cell lines respectively. Dashed lines indicate the mean of each distribution. The main cell line is in color while the other two conditions are overlaid in grey. The histograms are weighted such that each biological replicate contributes equally despite any differences in the number of technical replicates. **D-F)** Violin plots of per experimental measurements of mean, variance, and skew (respectively) of the zero-delay cross-correlation distribution. The color of each dot on the interior of the violin plot represents a different biological replicate. The overall plot is not weighted to account for this difference. **D)** The means of the zero-delay cross-correlations distributions were not statistically significant across cell lines(p = 0.363 for CTRL vs FAD, p = 0.554 for CTRL vs acAD, p = 0.897 for FAD vs acAD). Statistical significance is indicated by * (* p < 0.05, ** p < 0.01, *** p < 0.001).

Finally, we measured significant differences across combined distributions for the distance-limited cross correlation, although with only moderate effect sizes (KS (n = 5,545,197 cell pairs), Table S4). Similar to the zero-delay cross-correlation, the healthy CTRL distribution exhibits a secondary population of cell pairs at high cross-correlation coefficients, implying strong coordination across slow time-scales (Fig 5A). Interestingly, the FAD distribution also has a secondary population of cell pairs but at a different location with a lower cross-correlation value, representing some disruption of slow communication (Fig 5C). The acAD distribution displays a stronger tail, but not a secondary peak such as the CTRL or FAD distributions (Fig 5B). The acAD distribution lacks the strong population of highly-correlated cell pairs, but the reduction in slow communication is somewhat mitigated.

**Fig. 5.**
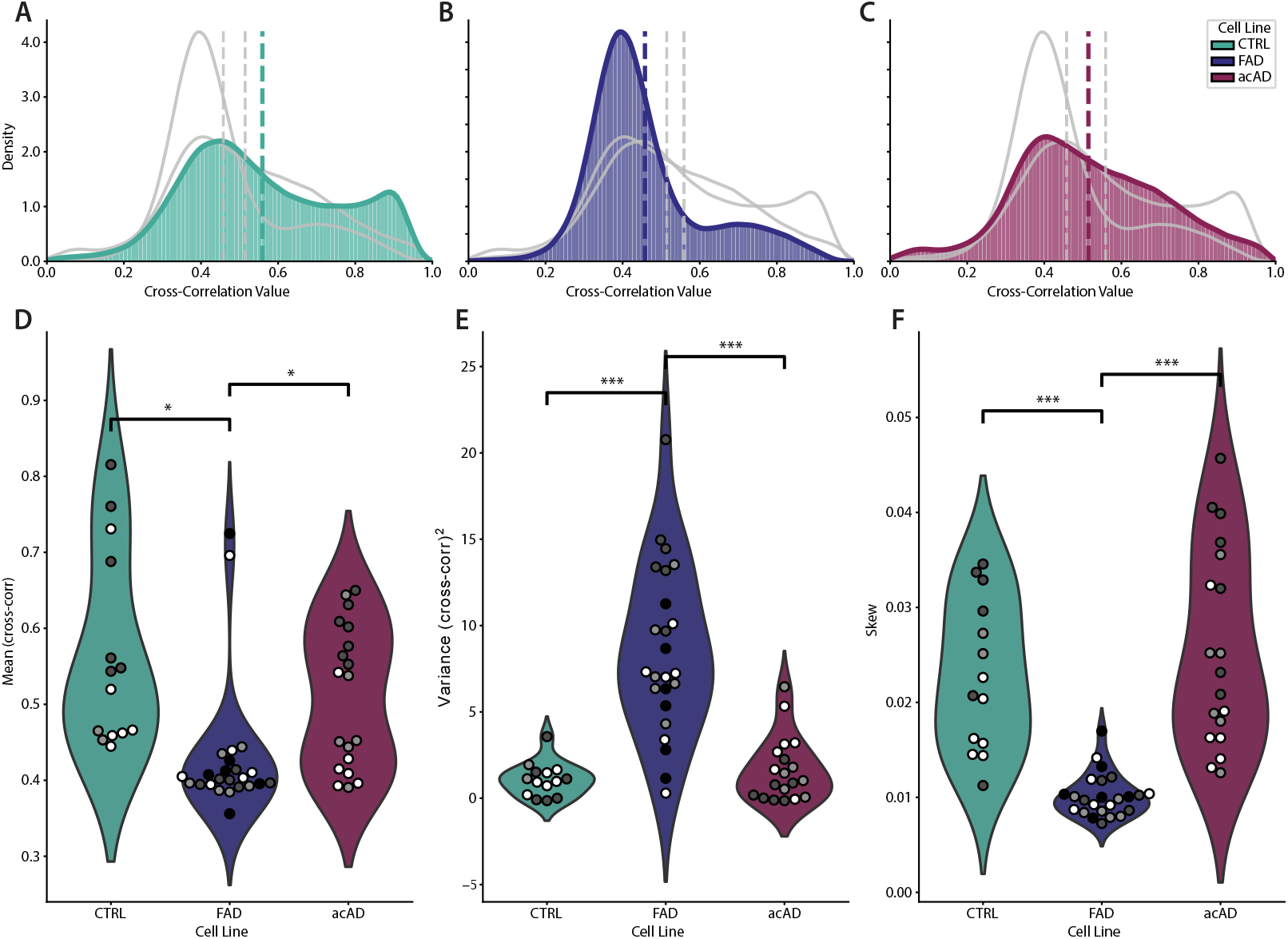
Characterization of distance-limited cross-correlation distributions of healthy and mutated cell lines. **A-C)** Density histograms of distance-limited cross-correlation coefficients for all cells identified across all experimental replicates for CTRL, FAD, and acAD cell lines respectively. Dashed lines indicate the mean of each distribution. The main cell line is in color while the other two conditions are overlaid in grey. The histograms are weighted such that each biological replicate contributes equally despite any differences in the number of technical replicates. **D-F)** Violin plots of per experimental measurements of mean, variance, and skew (respectively) of the distance-limited cross-correlation distribution. The color of each dot on the interior of the violin plot represents a different biological replicate. The overall plot is not weighted to account for this difference. Statistical significance is indicated by * (* p < 0.05, ** p < 0.01, *** p < 0.001).

The shapes of the distance-limited cross-correlation on a per-experimental basis correspond to differences observed in the combined distributions. Once again, KW tests confirm that there is a distinct effect due to cell line for all three moments (n = 57 recordings, Table S7). Pairwise tests of the mean, variance, and skew indicate strong differences in the slow communication between the CTRL and FAD networks as well as between the FAD and acAD networks (Fig 5D-F). The FAD networks display a lower mean than the CTRL and acAD networks, suggesting that there is a loss of coordination on slower time-scales. Additionally, the lower skew in FAD networks reflects a loss of high-correlation cell pairs, affirming an overall disruption of slow communication. This may be due to an impact on astrocytes and astrocytic communication in the networks, or other methods of neuronal communication such as the release of neuropeptides. Consistent with other metrics, the acAD networks do not show the same features of network dysfunction.

## Discussion

Here we carried out systematic analysis of spontaneous neural network activity, contrasting an established Alzheimer’s model FAD with our accelerated model acAD. We find that the two models have complementary features and strengths that could be harnessed to elucidate early functional impacts of AD on neural network information processing, learning, and memory.

In the FAD model we were able to show early changes in the internal state of the network, as eluicidated from spontaneous neural activity, even before it has been shown to display other biomarkers of AD such as Aβ and aggregated p-tau. Choi et al. reported Aβ plaques and elevated levels of Aβ and p-tau after six weeks of differentiation (40). We measured networks after three weeks of differentiation, when the acAD model displays Aβ plaques (26). The FAD model displays a larger fraction of hyperactive cells, which agrees with other findings that the average rate of calcium transients was unaltered in the same FAD model, but they observed an increase in the proportion of hyperactive cells (11). FAD networks also displayed a diminished tail of highly correlated pairs of cells, both in instantaneous and delayed correlations, indicating a loss of coordination between cells.

The acAD model exhibits complementary characteristics: despite the addition of multiple disease mutations, and the early emergence of structural characteristics of AD, the acAD model preserves spontaneous calcium activity and network organization. The comparable baseline for spontaneous calcium activity in the acAD and control networks lends itself to future studies of learning. One limitation of this study is that the observed calcium activity of the cells is spontaneous rather than evoked, which is often considered “noise” in animal models. In order to better understand AD and the acAD model, future work should focus on stimulated activity, specifically in a learning paradigm. Assessing effects of FAD mutations on signaling during and after learning may provide mechanistic insights. Due to the similar baseline of “noise correlations” in the acAD and the control, it is likely possible to disentangle differences in learning due to contributions from synaptic plasticity alone. Deviations observed between the control and the acAD model can be attributed to downstream biomarkers of AD, such as Aβ and p-tau, whereas deviations between the acAD model and the FAD model can be attributed to differences in the internal network state.

One limitation of this current study is that the calcium dynamics captured cannot be directly separated by cell type. The calcium dye we used is not specific to astrocytes or neurons, so the contributions of each cell type was not directly observed. While we attempt to infer differences based on the zero-delay and distance-limited cross-correlations, further work would be required to distinguish specific changes in neuronal and astrocytic signaling in these AD models. Previous research in a mouse model with similar FAD mutations (APP Swedish (KM670/671NL), PSEN1 with the L166P mutation) observed hyperactivity in astrocyte calcium signals that preceded neuronal hyperactivity (36). It is possible that this phenotype is present in the acAD or FAD models as well. Based on our findings, a natural question arises: why does FAD alone appear more functionally disrupted than the acAD model which includes an additional disease mutation? The acAD model may induce more stress on the neural cells, altering network maturation trajectories. Oxidative stress, or the generation of excessive reactive oxygen species (ROS), is a hallmark of not only aging in the central nervous system, but also with neurodegenerative disorders specifically (41). These ROS molecules can then trigger disruptions in Ca^2+^ homeostasis. One such example is the mutation to PSEN1 in the acAD model that is described above, which impacts the regulation of calcium ions in the endoplasmic reticulum. Recent work from our group found that the changes in ROS driven by UV light impact calcium signaling in our healthy hNPCs (42). This indicates that future work should explore the impact of ROS perturbations in both FAD and acAD models.

Overall, the FAD model exhibited early network dysfunction and a hyperactive subpopulation. Interestingly, despite rapidly developing hallmark AD-associated pathology, the acAD model retained spontaneous activity patterns more similar to healthy controls than to FAD networks. This divergence may indicate that structural pathology and spontaneous network dysfunction arise through partially distinct biological mechanisms or temporal trajectories. Such separation provides an opportunity in future work to experimentally disentangle pathological burden from emergent network-level dysfunction in human neural systems.

Taken together, these results suggest that pathological biochemical burden and dysfunction of the spontaneous network activity do not evolve synchronously and that their role in dysfunctions of information processing can be investigated separately. This highlights a new line of questioning for AD research.

## ACKNOWLEDGEMENTS

This material is based upon work supported by the Air Force Office of Scientific Research Biophysics Program under Award No. FA9550-25-1-0002. We would like to thank Sylvester J. Gates III for initial visualization adapted for figure 1. We would also like to thank Sylvester J. Gates III and K. Jeneh Perry for discussions and their biological expertise. Finally, we thank Corey Herr, Hoony Kang, Noah Chongsiriwatana, and Ian Whitehouse for helpful conversations around analysis methods.

## AUTHOR CONTRIBUTIONS

**Anna M. Emenheiser:** Conceptualization, Methodology, Imaging, Software, Visualization, Formal analysis, Writing - Original draft preparation. **Emma Gentry:** Conceptualization, Cell culture, Writing - review & editing. **Huijing Xue:** Conceptualization, Methodology, Cell culture, Writing - review & editing. **Phillip Alvarez:** Microscope design & instrumentation. **Kate M. O’Neill:** Visualization, Supervision, Writing - review & editing. **Kan Cao:** Conceptualization, Resources, Supervision, Funding acquisition, Writing - review & editing. **Wolfgang Losert:** Conceptualization, Resources, Supervision, Funding acquisition, Writing - review & editing.

## Notes

### Competing Interest Statement

The authors have declared no competing interest.

