## Supplemental Tables and Figures for "Network-Level Characterization of Spontaneous Calcium Activity in an In-Vitro Alzheimer’s Disease Model"

### Supplementary Note 1: Additional Data

Additional data was taken at a week 2 timepoint before even the acAD model is in the AD-like state. Analysis and visualization utilized the same methods as week 3, but not included in the main manuscript for brevity.

| Cell Line | Alzheimer's Mutations | Progerin Mutation | Week 2 Biological Replicates | Week 2 Technical Replicates | Week 3 Biological Replicates | Week 3 Technical Replicates |
|---|---|---|---|---|---|---|
| Non-transduced (CTRL) | | | 4 | 4; 5; 6; 6 | 3 | 2; 6; 6 |
| GFP control | | | 4 | 5; 6; 4; 6 | 3 | 3; 6; 6 |
| mCherry control | | | 2 | 6; 6 | 2 | 6; 6 |
| Lamin A (LA) | | | 4 | 6; 6; 1; 6 | 3 | 5; 5; 6 |
| Progerin (Pg) | | X | 4 | 6; 6; 5; 6 | 3 | 3; 6; 6 |
| mCherry + LA | | | 3 | 6; 6; 6 | 4 | 6; 6; 6; 6 |
| mCherry + Pg | | X | 3 | 6; 6; 6 | 3 | 6; 3; 6 |
| FAD | X | | 4 | 6; 6; 6; 6 | 4 | 6; 6; 6; 6 |
| FAD + LA | X | | 2 | 6; 4 | 3 | 6; 6; 6 |
| FAD + Pg (acAD) | X | X | 2 | 4; 6 | 3 | 7; 6; 6 |

**Table S1.** Description of full dataset collected, including number of biological and technical replicates collected at week two and week three. The technical replicates are split based on the number of technical replicates collected per biological replicate.

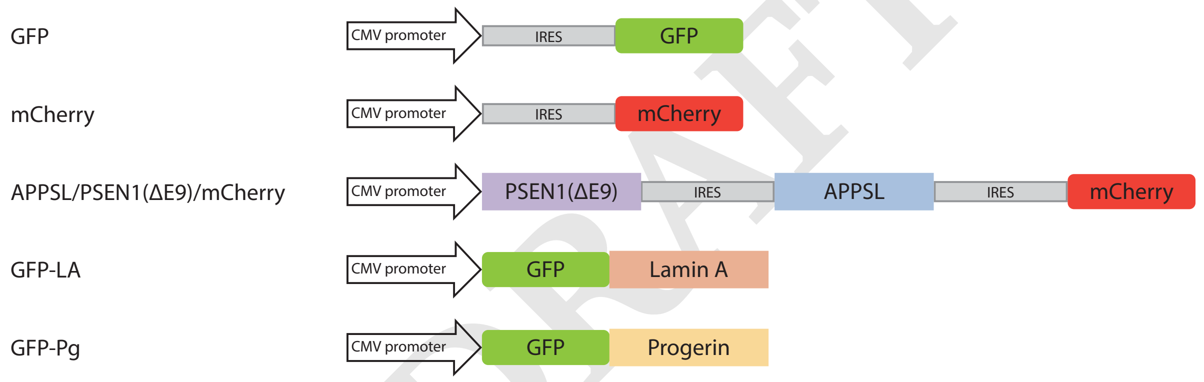


**Fig. S1. Genetic constructs used to create the different cell lines.** GFP was used for the GFP control. mCherry was used for the mCherry control. GFP-LA was used for the Lamin A (LA) cell line. GFP-Pg was used for the Progerin (Pg) cell line. A combination of mCherry and GFP-LA was used for mCherry + LA. A combination of mCherry and GFP-Pg was used for mCherry + Pg. APPSL/PSEN1ΔE9/mCherry was used for the FAD model (24). A combination of APPSL/PSEN1ΔE9/mCherry and GFP-LA was used for FAD + LA. Finally, a combination of APPSL/PSEN1ΔE9/mCherry and GFP-Pg makes up the acAD model (26).

DRAFT

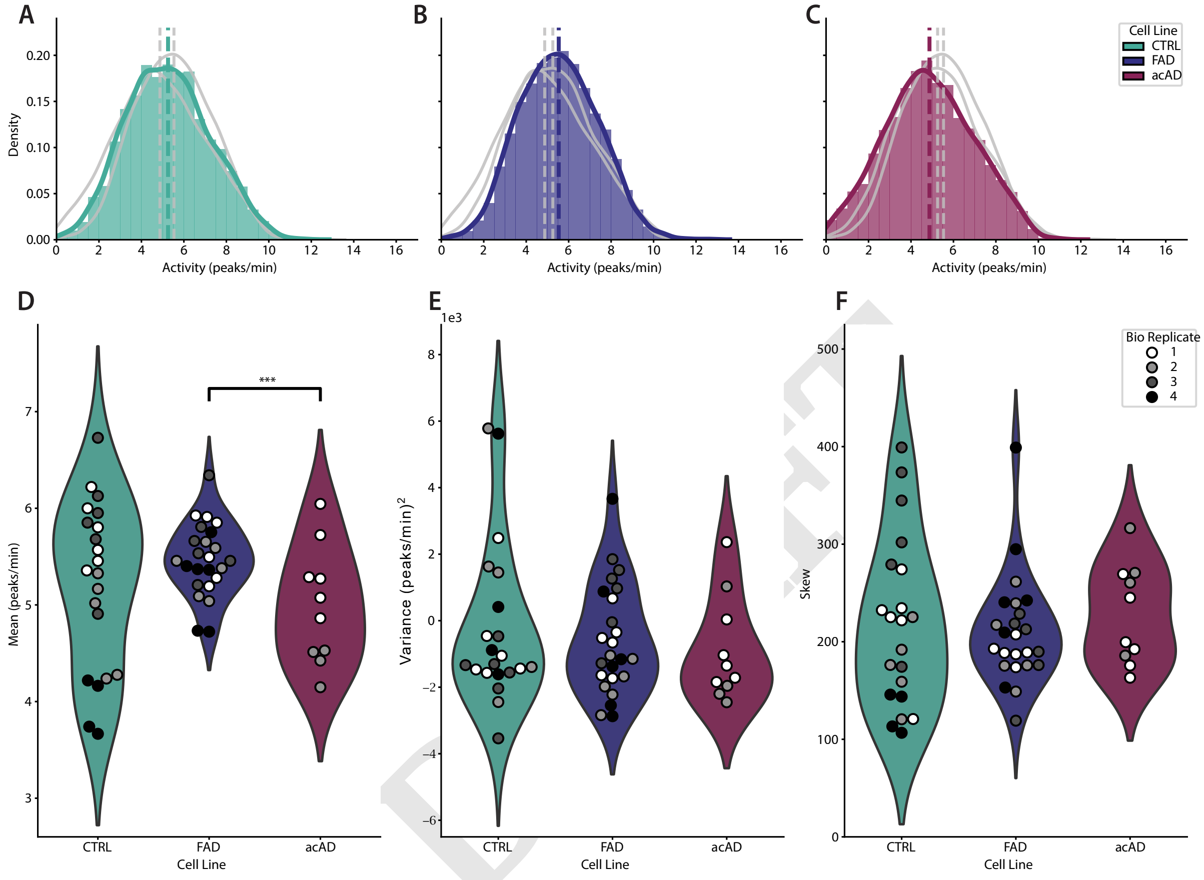


**Fig. S2. Characterization of calcium activity (spikes/min) of healthy and mutated cell lines at week two. A-C)** Density histograms of calcium activity (spikes/min) for all cells identified across all experimental replicates for CTRL, FAD, and acAD cell lines respectively. Dashed lines indicate the mean of each distribution. The main cell line is in color while the other two conditions are overlaid in grey. The histograms are weighted such that each biological replicate contributes equally despite any differences in the number of technical replicates.**D-F)** Violin plots of per experimental measurements of mean, variance, and skew (respectively) of the activity distribution. The color of each dot on the interior of the violin plot represents a different biological replicate. The overall plot is not weighted to account for this difference. Statistical significance is indicated by * (* $p < 0.05$, ** $p < 0.01$, *** $p < 0.001$).

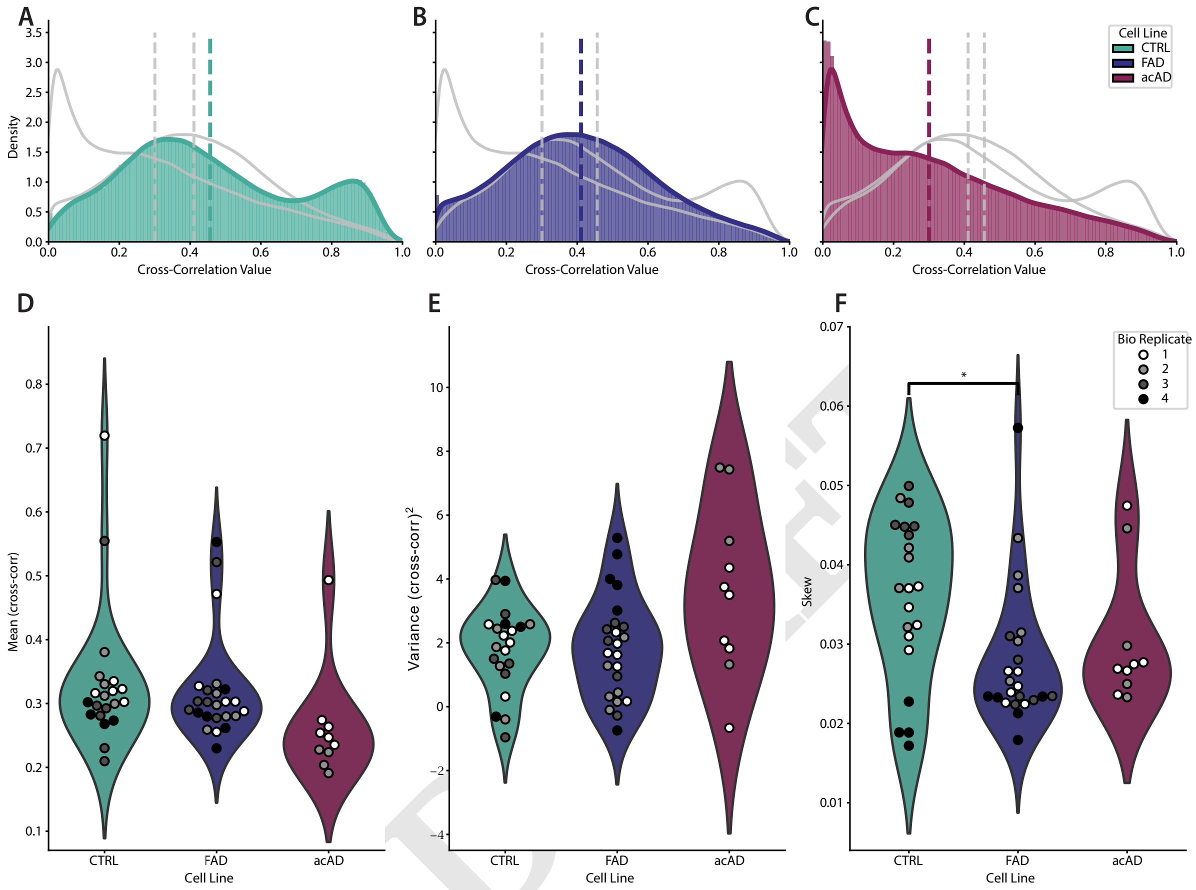


**Fig. S3. Characterization of zero-delay cross-correlation distributions of healthy and mutated cell lines at week two. A-C)** Density histograms of zero-delay cross-correlation coefficients for all cells identified across all experimental replicates for CTRL, FAD, and acAD cell lines respectively. Dashed lines indicate the mean of each distribution. The main cell line is in color while the other two conditions are overlaid in grey. The histograms are weighted such that each biological replicate contributes equally despite any differences in the number of technical replicates. **D-F)** Violin plots of per experimental measurements of mean, variance, and skew (respectively) of the zero-delay cross-correlation distribution. The color of each dot on the interior of the violin plot represents a different biological replicate. The overall plot is not weighted to account for this difference. **D)** The means of the zero-delay cross-correlations distributions were not statistically significant across cell lines($p = 0.363$ for CTRL vs FAD, $p = 0.554$ for CTRL vs acAD, $p = 0.897$ for FAD vs acAD). Statistical significance is indicated by * (* $p < 0.05$, ** $p < 0.01$, *** $p < 0.001$).

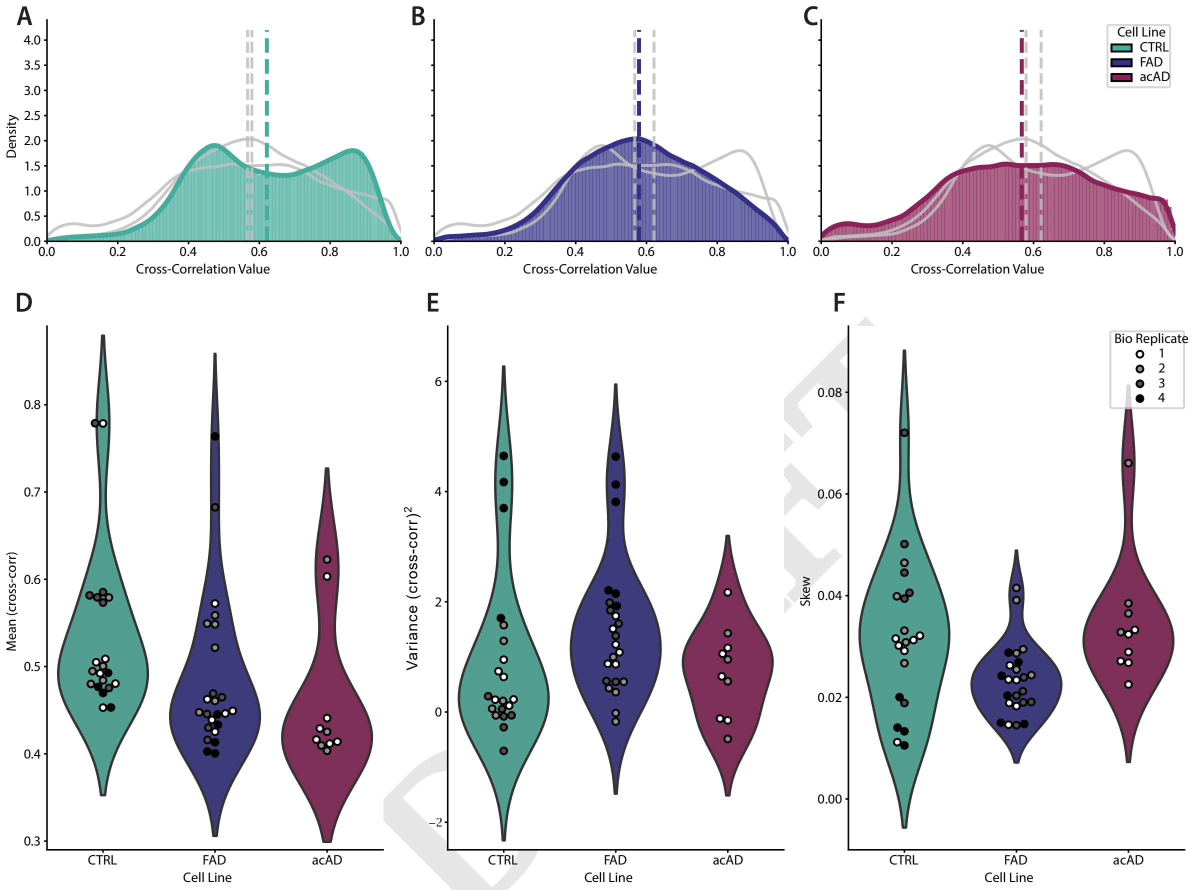


**Fig. S4. Characterization of distance-limited cross-correlation distributions of healthy and mutated cell lines at week two. A-C)** Density histograms of distance-limited cross-correlation coefficients for all cells identified across all experimental replicates for CTRL, FAD, and acAD cell lines respectively. Dashed lines indicate the mean of each distribution. The main cell line is in color while the other two conditions are overlaid in grey. The histograms are weighted such that each biological replicate contributes equally despite any differences in the number of technical replicates. **D-F)** Violin plots of per experimental measurements of mean, variance, and skew (respectively) of the distance-limited cross-correlation distribution. The color of each dot on the interior of the violin plot represents a different biological replicate. The overall plot is not weighted to account for this difference. Statistical significance is indicated by * (* $p < 0.05$, ** $p < 0.01$, *** $p < 0.001$).

### Supplementary Note 2: Statistical Tests

| Cell Line A | Cell Line B | Kolmogorov–Smirnov Stat | Corrected p-value |
|---|---|---|---|
| CTRL | FAD | 0.1513 | $< 1 \times 10^{-16}$ |
| CTRL | acAD | 0.0648 | $< 1 \times 10^{-16}$ |
| FAD | acAD | 0.1914 | $< 1 \times 10^{-16}$ |

**Table S2.** Complete Kolmogorov–Smirnov pairwise comparison with Holm-Bonferroni correction between combined distributions of activity (spikes/min) at week three.

| Cell Line A | Cell Line B | Kolmogorov–Smirnov Stat | Corrected p-value |
|---|---|---|---|
| CTRL | FAD | 0.1443 | $< 1 \times 10^{-16}$ |
| CTRL | acAD | 0.0929 | $< 1 \times 10^{-16}$ |
| FAD | acAD | 0.1307 | $< 1 \times 10^{-16}$ |

**Table S3.** Complete Kolmogorov–Smirnov pairwise comparison with Holm-Bonferroni correction between combined distributions of zero-delay cross-correlation at week three.

| Cell Line A | Cell Line B | Kolmogorov–Smirnov Stat | Corrected p-value |
|---|---|---|---|
| CTRL | FAD | 0.2941 | $< 1 \times 10^{-16}$ |
| CTRL | acAD | 0.0924 | $< 1 \times 10^{-16}$ |
| FAD | acAD | 0.2300 | $< 1 \times 10^{-16}$ |

**Table S4.** Complete Kolmogorov–Smirnov pairwise comparison with Holm-Bonferroni correction between combined distributions of distance-limited cross-correlation at week three.

| Metric | Kruskal-Wallis Stat | p-value |
|---|---|---|
| Mean | 19.4192 | $6.0697 \times 10^{-5}$ |
| Variance | 16.7307 | $2.3279 \times 10^{-4}$ |
| Skew | 25.8320 | $2.4584 \times 10^{-6}$ |

**Table S5.** Complete Kruskal-Wallis tests for each moment of per-experimental distributions of activity (spikes/min) at week three.

DRAFT

| Metric | Kruskal-Wallis Stat | p-value |
|---|---|---|
| Mean | 1.9942 | $3.6894 \times 10^{-1}$ |
| Variance | 25.6964 | $< 1 \times 10^{-6}$ |
| Skew | 31.6372 | $< 1 \times 10^{-7}$ |

**Table S6.** Complete Kruskal-Wallis tests for each moment of per-experimental distributions of zero-delay cross-correlation at week three.

DRAFT

| Metric | Kruskal-Wallis Stat | p-value |
|---|---|---|
| Mean | 21.3310 | $2.3336 \times 10^{-5}$ |
| Variance | 31.4072 | $1.5136 \times 10^{-7}$ |
| Skew | 37.1405 | $8.6107 \times 10^{-9}$ |

**Table S7.** Complete Kruskal-Wallis tests for each moment of per-experimental distributions of distance-limited cross-correlation at week three.

### Supplementary Note 3: Extended Analysis Methods

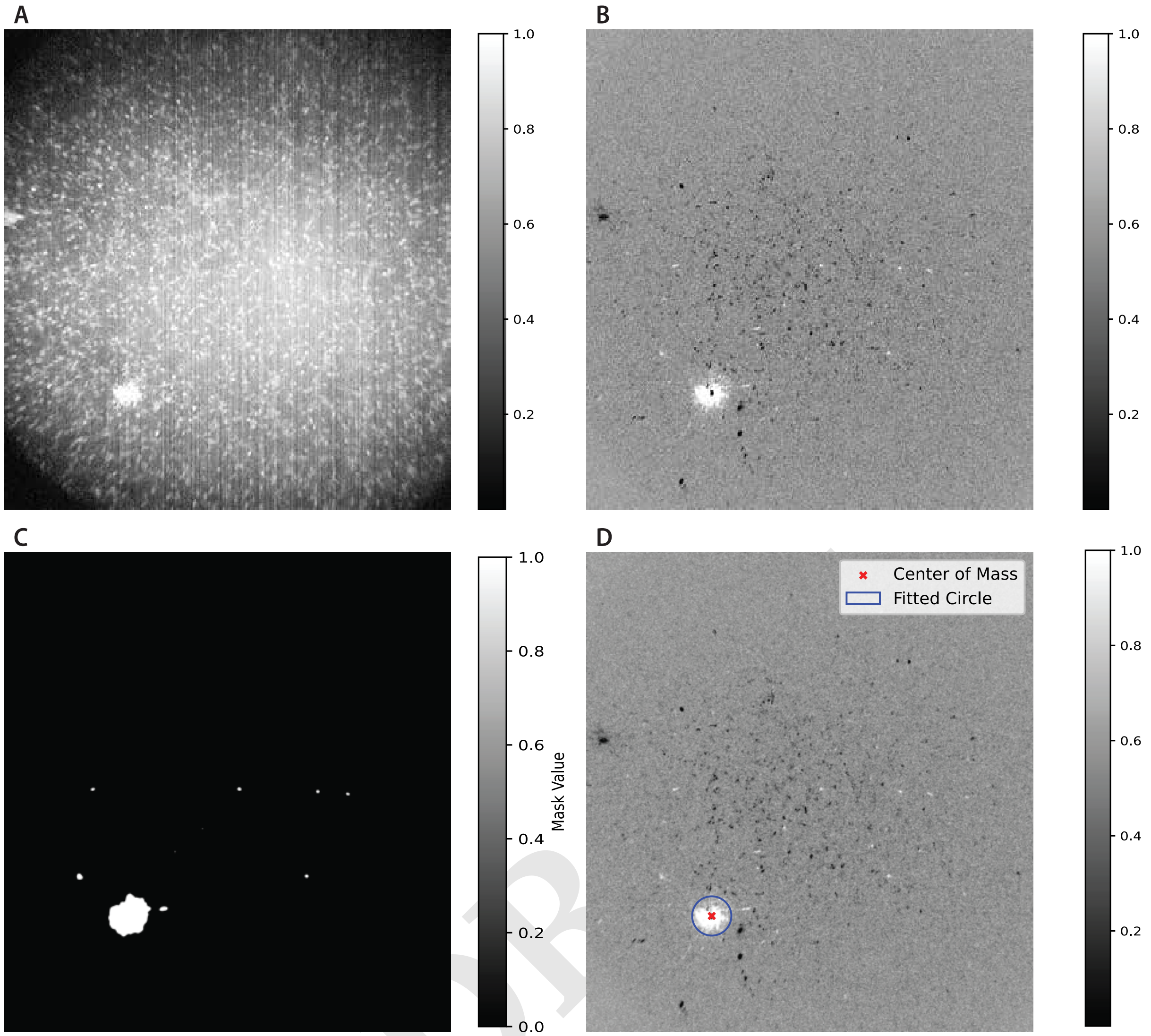


**Fig. S5. Workflow to calculate the speed of calcium flow in hNPCs. A)** A standard deviation projection of frames corresponding to a localized burst of calcium activity in a control network recording. **B)** A difference image of the standard deviation projection for the frames of the burst and a median projection of the entire movie. **C)** A binary mask created by thresholding the difference image. A filter was applied in order to remove small objects from the binary mask. **D)** A fitted circle centered on the center of mass of the binary mask. The final speed of calcium flow was estimated based on the radius of the circle and the time between the start and end frames of the burst event. This was done for n = 10 bursts observed across N = 5 movies.
